# Establishment of signaling interactions with cellular resolution for every cell cycle of embryogenesis

**DOI:** 10.1101/285007

**Authors:** Long Chen, Vincy Wing Sze Ho, Ming-Kin Wong, Xiaotai Huang, Lu-yan Chan, Hon Chun Kaoru Ng, Xiaoliang Ren, Hong Yan, Zhongying Zhao

## Abstract

Intercellular signaling interaction plays a key role in breaking fate symmetry during animal development. Identification of the signaling interaction at cellular resolution is technically challenging, especially in a developing embryo. Here we develop a platform that allows automated inference and validation of signaling interaction for every cell cycle of *C. elegans* embryogenesis. This is achieved by generation of a systems-level cell contact map that consists of 1,114 highly confident intercellular contacts by modeling analysis and is validated through cell membrane labeling coupled with cell lineage analysis. We apply the map to identify cell pairs between which a Notch signaling interaction takes place. By generating expression patterns for two ligands and two receptors of Notch signaling pathway with cellular resolution using automated expression profiling technique, we are able to refine existing and identify novel Notch interactions during *C. elegans* embryogenesis. Targeted cell ablation followed by cell lineage analysis demonstrates the roles of signaling interactions over cell division in breaking fate symmetry. We finally develop a website that allows online access to the cell-cell contact map for mapping of other signaling interaction in the community. The platform can be adapted to establish cellular interaction from any other signaling pathways.

## Introduction

Symmetry breaking in cell division timing and cell fate specification has long been a focus of developmental biology. Intercellular signaling plays a key role in breaking these symmetries (Yochem *et al*. 1988; Sawa 2012; Clevers and Nusse 2012; Greenwald 2013; Zacharias *et al*. 2015) although maternal control is critical for establishing polarity during early development (Rose and Gonczy 2014). For example, a Notch signaling interaction is necessary for fate asymmetry between cells ABa and ABp (Mickey *et al*. 1996; Priess 2005); whereas a Wnt interaction is required for both fate asymmetry and division asynchrony between cells EMS and P2 in a four-cell *Caenorhabditis elegans* embryo(Rocheleau *et al*. 1997). The Notch interaction is achieved by a contact between the P2 that expresses a Notch ligand, *apx-1*, and the ABp but not the ABa cell, although both the later cells express Notch receptor, *glp-1* (Mickey *et al*. 1996). This demonstrates that a contact between cells is essential for triggering a signaling interaction to drive differential fate specification (Good *et al*. 2004). A similar scenario is observed for the Wnt interaction between the EMS and the P2 cells, which is necessary for asymmetric division of the former into MS and E cells during *C. elegans* embryogenesis (Goldstein 1992; Rocheleau *et al*. 1997). Notably, the two pathways are used repeatedly throughout development in a cellular context-dependent fashion to establish further asymmetries in fate specification or division timing (Huang *et al*. 2007; Zacharias *et al*. 2015). For example, in a 12-cell *C. elegans* embryo, the four great-granddaughters of AB express Notch receptor, GLP-1, but only two of them, i.e., ABalp and ABara, are in contact with a Notch ligand-expressing cell, MS, leading to their differential differentiation into mesodermal and ectodermal fates, respectively (Hutter and Schnabel 1994; Shelton and Bowerman 1996). Importantly, signaling interactions from the same pathway may have an opposite consequence depending on their timing or cellular context. For example, the first Notch interaction inactivates its targets, *tbx-37/38* (Good *et al*. 2004); whereas the second one activates its targets including PHA-4, a FoxA transcription factor required for pharynx organogenesis (Priess 2005). These time-dependent signaling events indicate that dissecting signaling interactions with precise spatial and temporal resolution would be essential for a thorough understanding of symmetry breaking during metazoan development.

One of the biggest challenges in defining a signaling interaction during embryogenesis is the establishment of cell identity, especially in an embryo with a large number of cells (Keller *et al*. 2008; Zacharias and Murray 2016). Another challenge is that one must have access to the cellular expression patterns of signaling molecules for each cell cycle. These requirements inhibit functional characterization of cellular signaling during rapid development. This is because defining a signaling interaction requires knowledge on the identities of cell pairs that are in contact with each other, with one expressing a ligand and the other a receptor.

The development of cell-tracking techniques using time-lapse 3D (hereafter referred to as 4D) microscopy has greatly facilitated cell lineage analysis (Schnabel *et al*. 1997, 2006, Zhao *et al*. 2008, 2010b; Muzzey and van Oudenaarden 2009). In particular, a recently developed automated lineaging technique allows routine tracing of cell division and single-cell expression profiling in a *C. elegans* embryo up to 350 cells within approximately half an hour and up to the last round of cell division of embryogenesis in about one day (Bao *et al*. 2006; Murray *et al*. 2008; Richards *et al*. 2013; Du *et al*. 2014; Shah *et al*. 2017). This technique makes it possible to infer signaling interaction at cellular resolution for every cell cycle (Fig. 1) because the output of automated lineaging contains quantitative positional information for nuclei of all cells for every minute during embryogenesis, thus allowing systematic modelling of cell contacts with exceptional spatial and temporal resolution. A cell contact map up to the ∼150-cell stage was reported for the *C. elegans* embryo purely based on Voronoi modeling (Hench *et al*. 2009). However, the map suffers from several caveats. First, it was generated using a single “composite” embryo assembled from six different embryos, each of which was partially resolved for cell lineage. Given the variability in embryo size, shape, and developmental timing (Hara and Kimura 2009; Greenan *et al*. 2010; Moore *et al*. 2013; Ho *et al*. 2015), it would be problematic to superimpose the six embryos into a single embryo for modeling of cell contact. Second, a thorough validation of the modeling results was not performed. Many cell contacts that are brief in duration and/or have a minimal contact area may not be consequential. As a result, a relatively high false-positive rate is unavoidable without taking these issues into account. Finally, the map covers only the ∼150-cell stage, but a *C. elegans* embryo does not hatch until it develops into 558 cells (Sulston *et al*. 1983). Therefore, a more reliable cell contact map that covers cells born at a later stage of embryogenesis is necessary for dissecting cell signaling. Here, we present a platform that allows the automated inference of cellular signaling for every cell cycle up to the ∼350-cell *C. elegans* embryo. Applying the platform to Notch signaling pathway demonstrated a consecutive signaling events over cell cycles for breaking cell fate symmetry.

**Fig 1.**
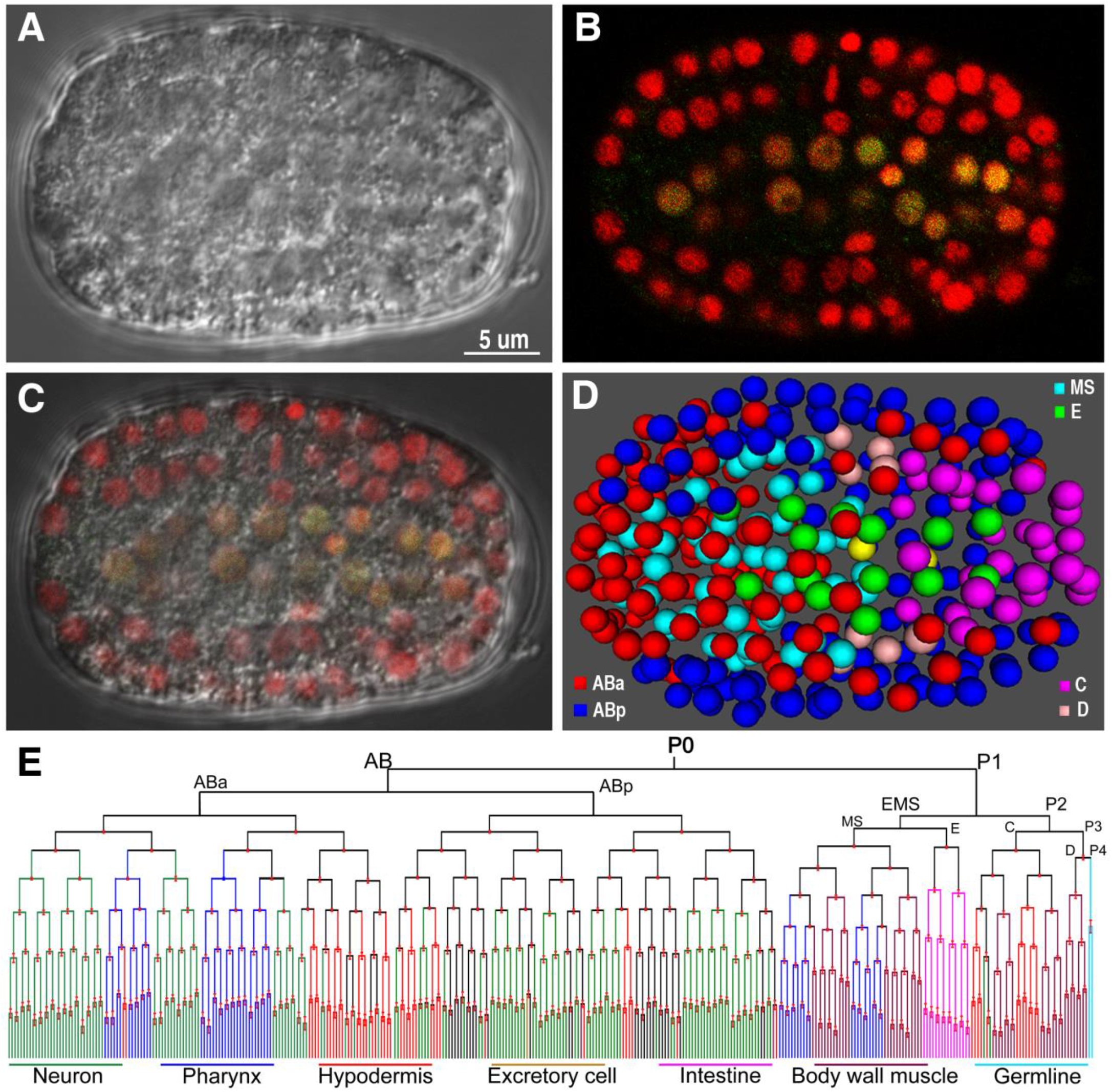
Overview of a 350-cell stage *C. elegans* embryo. A. Nomarski micrograph of a *C. elegans* embryo of appropriate 350-cell stage. B. Epifluorescence micrograph showing superimposed nuclear expression patterns of lineaging markers (red) and a pharynx marker, PHA-4 (green). C. Superimposed micrograph from panels B and C. D. 3D space-filling model of a 350-cell embryo. Cells are differentially color-coded based on their lineal origins. E. Cell lineage tree up to 350-cell stage. Cell lineages are differentially colored based on cell fate except the undifferentiated cell fates that are colored in black.

## Results

### A time-lapse cell-contact map from 4- to 350-cell *C. elegans* embryo

To facilitate the precise assignment of cell pairs between which a potentially functional signaling interaction takes place, we performed modeling analysis of cell-cell contact over the proliferative stage of *C. elegans* embryogenesis from 4 to 350 cells. Specifically, 4D coordinates from 91 wild-type embryos generated previously by automated lineaging (Ho *et al*. 2015) were individually used as an input for the Voronoi algorithm to model cell surfaces, from which the contacting area is computed between a cell pair (see Materials and Methods). Instead of using partial 4D coordinates from different embryos, as in a previous study (Hench *et al*. 2009), the 91 coordinate sets used here were each derived from single intact embryos, which minimizes the issues associated with normalization steps for cell size, embryo shape and developmental timing.

It is conceivable that many cell contacts may not be relevant to cell signaling due either to their short duration or small contact area. To increase the modeling accuracy, we adopted the following criteria to define an effective cell contact, which is referred to as cell contact hereafter for simplicity unless stated otherwise. First, a contact area is required to be at least 6.5% of the average cell surface areas of all cells present at the same time point (Fig. 2A). Second, this criterion must be satisfied for at least two consecutive time points (approximately 1.5 minutes per time point) (Fig. 2C, see details below). Third, these two criteria must be reproducible in at least 95% of the 91 wild-type embryos (i.e., in 87 of 91 embryos; Fig. 2B). As a result, we predicted a total of 1,114 cell contacts from the 4- to 350-cell stage (Table S1). The predicted contact areas were highly reproducible among the 91 embryos with a Pearson correlation coefficient (r) of at least 0.8 between any two independent embryos (Fig. 2D). The predicted cell contact can be readily validated via ubiquitous and simultaneous labeling of cell membranes and nuclei with resolved cell identities (Fig. 2E). We adopted the criterion of a 6.5% contact area based on the well-established 2^nd^ Notch interactions in the *C. elegans* embryo (Mickey *et al*. 1996). This interaction occurs between a Notch ligand-expressing cell MS and two Notch receptor-expressing cells ABalp and ABara in a 12-cell embryo, but not in their sisters (ABala and ABarp), which leads to specification of their pharyngeal fate. We first individually computed the contact areas between MS and each of the four AB descendants for the 91 wild-type embryos. Given the variability in contact area between the embryos, we next plotted the occurrence of the four contacts (any contact with a contacting area > 0) in the 91 embryos against the ratio of actual contact area relative to average cell surface areas of all cells present at the current time point. Occurrence distributions of both individual (Fig. S1) and aggregated (Fig. 2A) plots demonstrated a normal distribution. We observed a clear demarcation between cell pairs with (between MS and ABala or ABarp) and without (between MS and ABalp or ABara) a functional contact at a ratio of approximately 6.5% of the actual contact area relative to the average cell surface area of all cells at the current time point (Fig. 2A). We therefore used the ratio of 6.5% as a cutoff for defining an effective contact. Variability in actual cell contact was observed not only between MS and the four AB descendants, but also in other cells from 4-350 cells in the 91 embryos (Fig. 2B). Therefore, we require that only if a contact is reproducibly observed in 95% of all the 91 embryos, it can be defined as an effective contact. To further reduce our false-positive rate in calling an effective cell contact, we require a contact that lasts for at least two consecutive time points (approximately 3 minutes). We set this filter because our temporal resolution is 1.5 minutes, meaning that the duration of any contact shorter than this will be assigned as 1.5 minutes. This temporal requirement ensures that an effective cell contact lasts for at least 1.5 minutes. A previous study suggested the substantial effect of pressure applied to an embryo during imaging on the prediction of cell-cell contact (Hench *et al*. 2009). We tested the effect of such pressure by examining whether the hatching rates are similar between pressured (mounted) and unpressurized (unmounted) embryos (those laid freely on an NGM plate). If the hatching rates are comparable, after the hatched larvae grow up, whether their brood sizes are comparable. We found that all mounted and unmounted embryos with 25 each hatched, and the brood sizes are also comparable between the mounted and unmounted embryos (Fig. S2), suggesting that pressure applied on the embryos for mounting was unlikely to have affects the important cell contacts during *C. elegans* embryogenesis.

**Fig 2.**
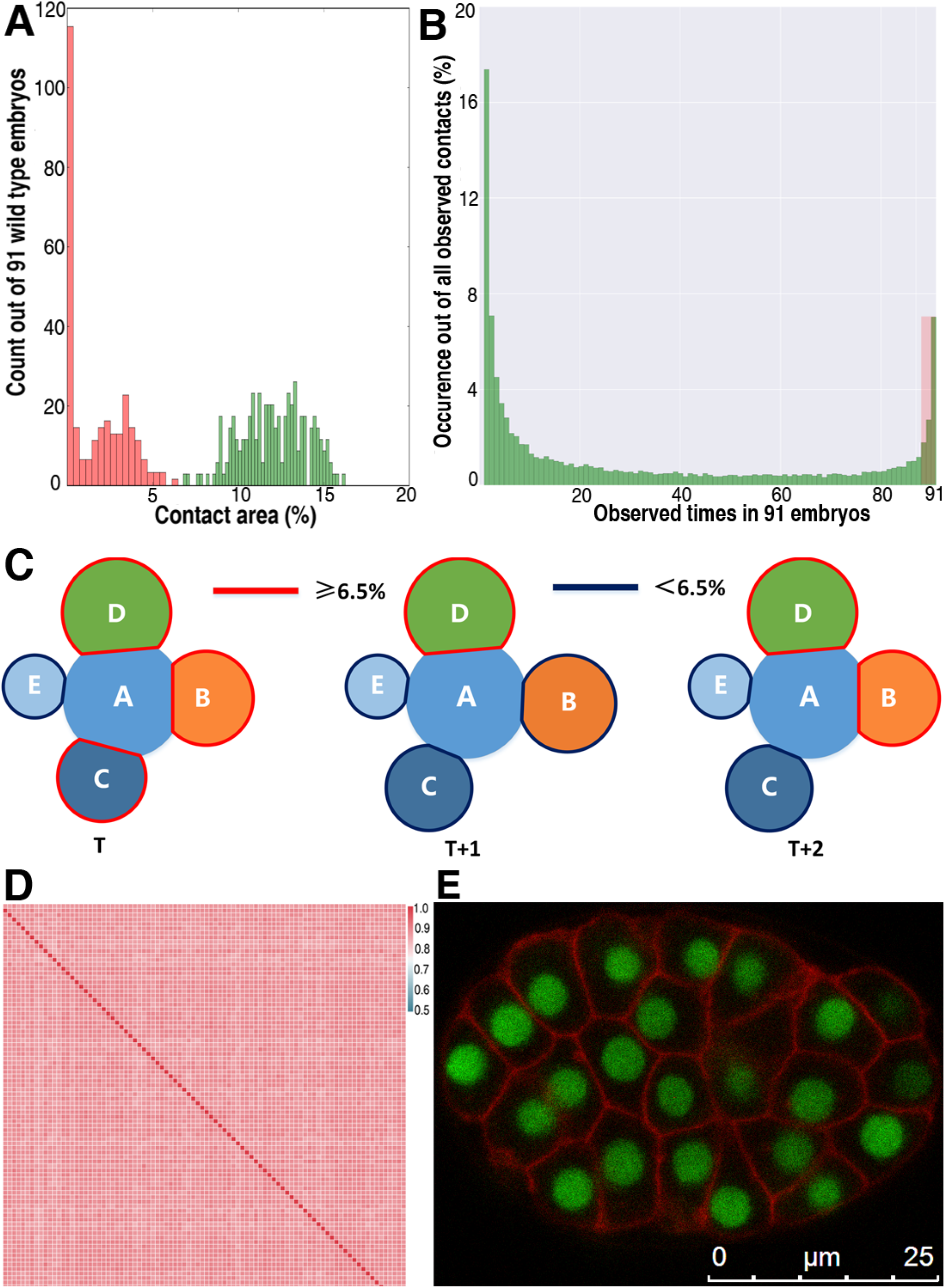
Modeling of cell-cell contact during *C. elegans* embryogenesis. A. Demarcation of the ratio of contact areas between cells with and without a functional contact. Shown is the occurrence distribution of modeled contact areas between cell pairs that are known to have (green) or not to have (brown) the 2^nd^ Notch (see main text) interactions in 91 embryos. Percentage of contact area out of average cell surface area of all cells in the current time point is plotted on x axis and the number of embryos with a given ratio out of 91 wild types on y axis. B. Distribution of any contacts (contact area >0) in 91 wild-type embryos. Y axis denotes the percentage of a given contact out of all observed contacts and x axis the observed times for a given contact out of 91 embryos. Contacts with over 95% reproducibility (i.e., observed in 87 out of 91 embryos) are shaded in red. C. A diagram showing the definition of effective cell contact with cell A over consecutive three time points. Contact area that is bigger or smaller than 6.5% of the average surface areas of all the cells is differentially colored in red and blue lines respectively. For a given cell pair, only cell contact area that is over 6.5% for at least two consecutive time point (around 3 minutes) is defined as an effective cell contact. D. Heat map of mutual Pearson correlation’s coefficient (*r*) of contact areas for all cells between 91 individual wild-type embryos. Both horizontal and vertical axes denote the coefficient of an individual embryo against another. E. An example of 40-cell *C. elegans* embryo expressing GFP in nuclei and membrane marker PH in cell membrane (red).

**Table 1.**
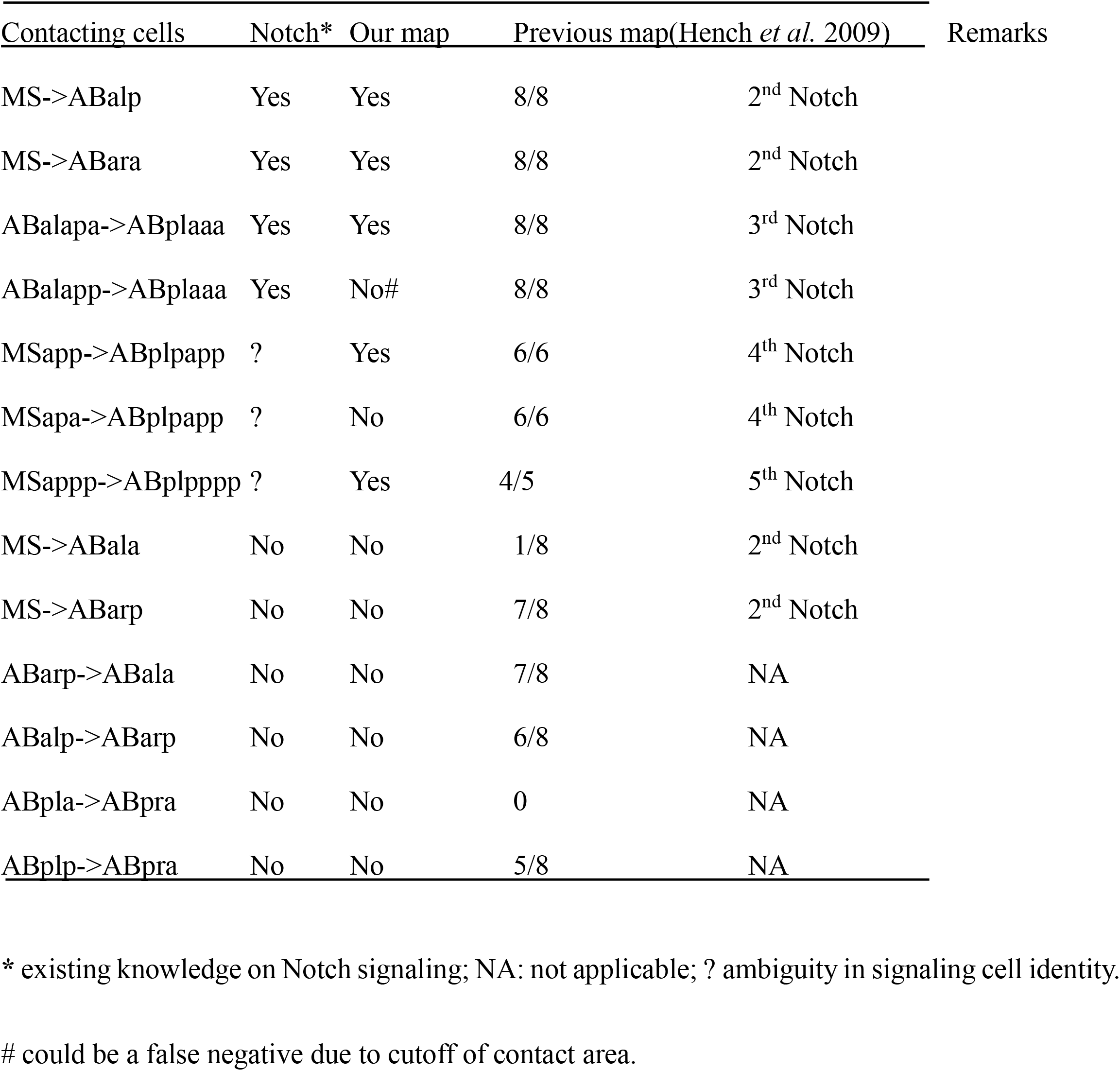
Comparison of our cell contact map with a previous map(Hench *et al*. 2009) and the contacting cell pairs with known Notch signaling interaction.

### Comparison of performance between our and a previous contact map

A previous cell contact map was generated with a modeling algorithm similar to that used here but using a single “composite” embryo assembled from six different embryos(Hench *et al*. 2009). The cell lineage for each embryo was partially resolved owing to the difficulty in establishing cell identities on both sides of an embryo. Notably, the cell contact was defined mainly based on whether there is any physical contact regardless of the size and duration of a contact, making it prone to a relatively high false-positive rate. The spatial and temporal constrains we used for modeling are expected to reduce the rate of false positives.

To compare the performances between our and the previous contact maps, we contrasted a subset of cell contacts relevant to well-established Notch signaling interactions (Table 1). It was expected that our modeling contacts would agree well with the contacts based on the 2^nd^ Notch interactions because they were used as a training set for our contact modeling. Notably, nearly one half of the cell contacts predicted previously were false positive when compared with the experimentally verified ones whereas our predictions agreed well with the experimental data from multiple Notch interactions (Table 1), indicating that our modeling method substantially outperforms the previous method in terms of accuracy.

### Lineal expression of Notch receptors and ligands derived from a single-copy transgene

Knowledge of the time-lapse expression of a ligand and its receptor of a signal pathway at the cellular level with high temporal resolution is critical for assigning a cell pair between which a signaling interaction takes place. However, such knowledge is either absent or present at poor spatiotemporal resolution especially during the proliferative stage of embryogenesis, thus preventing effective assignment of a signaling interaction. For example, existing expression patterns on Notch pathway components in *C. elegans* were obtained through either a transgenic study or antibody staining or their combination (Mello *et al*. 1994; Mickey *et al*. 1996; Moskowitz and Rothman 1996). Most of the transgenic assays are based on extrachromosomal arrays (Moskowitz and Rothman 1996) or biolistic bombardment (Murray *et al*. 2012). The expression patterns generated from these transgenic strains may suffer from increased perdurance of fluorescent reporter by extra copy of transgenes or uncertainty in regulatory sequences incorporated into host cells.

To generate the embryonic expression pattern of a Notch component that more likely mimics its native expression at cellular resolution for each cell cycle, we first produced multiple independent transgenic strains carrying a single copy of a fusion between GFP and a promoter sequence from a Notch component using the miniMos technique (Frokjaer-Jensen *et al*. 2014), including two functionally redundant receptors, *lin-12* and *glp-1*, and two ligands, *apx-1* and *lag-2*. A single strain that showed consistent expression with at least one another transgenic copy was used to map the reporter’s lineal expression using automated lineaging and expression profiling technology (Murray *et al*. 2008). *glp-1* shows specific expression in the descendants of ABarpap and ABplaaa (Fig. 3A, B, G and J). These will generate hypodermal cells found in the head (Sulston *et al*. 1983). Dim expression was also observed in the descendants of MSaa and MSpa (Fig. S2A, F). Notably, our expression patterns are roughly comparable with those derived from the transgenic strains generated with biolistic bombardment in AB (Murray *et al*. 2012), but expression was observed in more cells in the bombardment strains. Because the promoter sequences are similar in size, it remains likely that the expression conferred by the single-copy transgene may be too dim to be seen. Expression of the other Notch receptor, *lin-12*, is mainly observed in the descendants of ABplp, ABprp and ABplaaa (Fig. 3C, D, H, J). No expression was observed in the P1 sublineage (Fig. S3B, G). One Notch ligand, *apx-1*, showed expression mainly in the descendants of ABala, ABpl(r)apaa (Fig. 3E, F, I, J), MSppapp and MSppppp (Fig. S3C, H). We did not observe the expression of *lag-2* in the ABalap descendants, as reported previously (Moskowitz and Rothman 1996). A complete list of cell expressing Notch ligands and receptors are shown in Table S2. When combined with the cell contact map, the lineal expression of these Notch components at a 1.5-minute interval over development will not only allow validation of existing Notch signaling interactions, especially at a stage with tens to hundreds of cells, but it also holds promise for the identification of novel cell pairs between which a signaling interaction may take place. We illustrate the applications in detail below.

**Fig 3.**
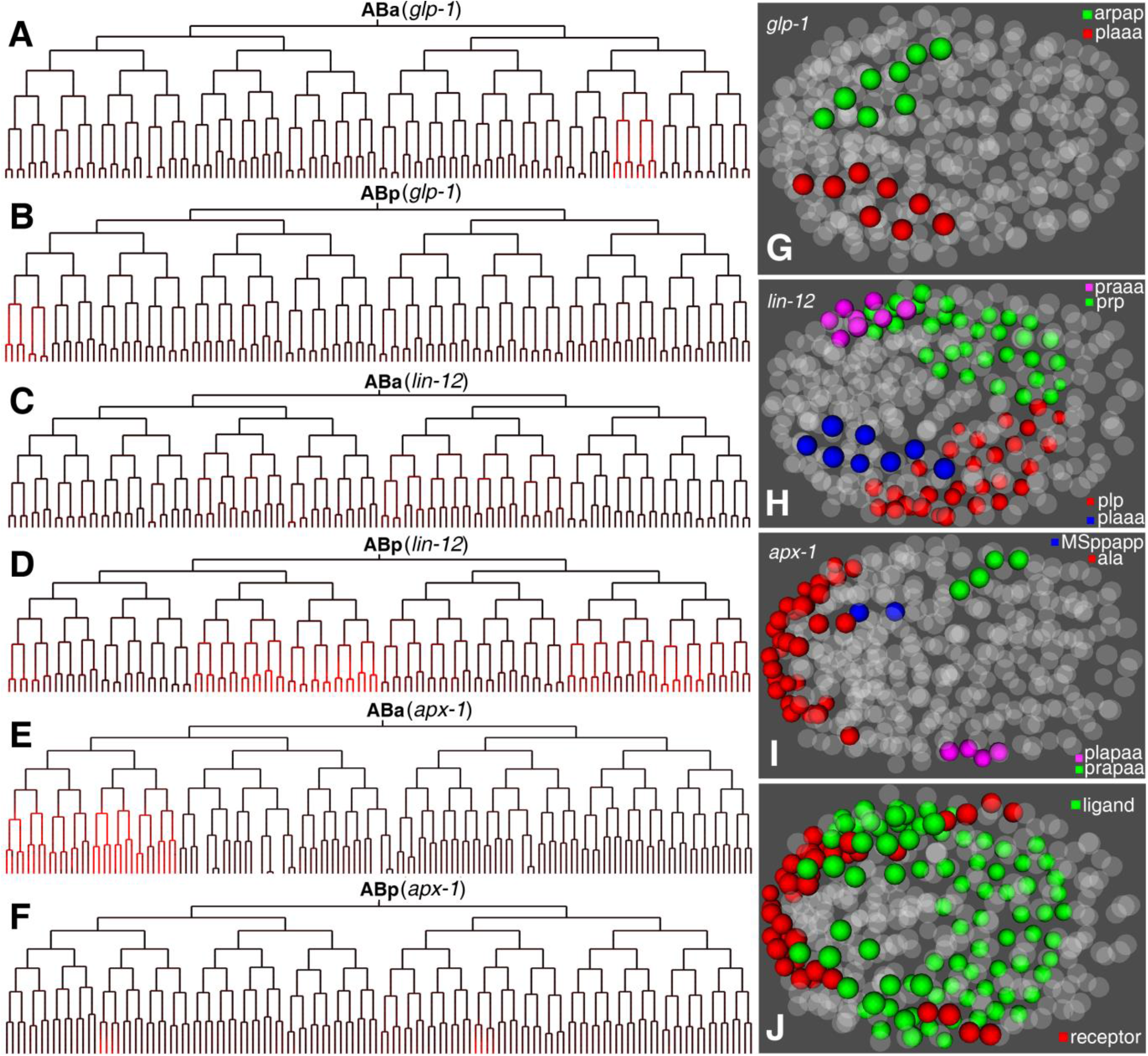
Expression of Notch ligands and receptors in *C. elegans* embryo. A-F. Lineal expression (red) of two Notch receptors, *glp-1* and *lin-12*, and one ligand, *apx-1*, in ABa and ABp lineage up to 350-cell stage. G-I. Spatial expression of the three genes differentially color coded based on their lineal origins. J. Combined spatial expression patterns of the three genes between ligand, *apx-1*, (red) and the two receptors, *glp-1* and *lin-12* (green).

### Refinement of the proposed cell pairs for 3^rd^ Notch interaction in *C. elegans* embryo

The 3^rd^ Notch signaling between signaling cell ABalapp and signal-receiving cell ABplaaa was proposed mainly based on the expression timing of a Notch ligand, *lag-2*, in ABalapp and a Notch receptor, *lin-12*, in the ABplaaa (Moskowitz and Rothman 1996). To confirm the signaling interaction and examine its functional redundancy, we took advantage of our time-lapse cellular expression patterns of both Notch receptors and two different ligands and aligned them against our modeled cell contacts. If a cell contact is observed between two cells with one expressing a ligand and the other a receptor, it is plausible that a signaling interaction takes place between the two. In addition to *lag-2* (Moskowitz and Rothman 1996), we observed the expression of another Notch ligand, *apx-1*, in the descendants of ABala (Fig. 3E). Our reporter assay revealed that both Notch receptors, *lin-12* and *glp-1*, are expressed in the left head precursor, ABplaaa (Fig. 3B, D). Despite the expression of *apx-1* in all of the descendants of ABala (Fig. 3E), only one of the ABala daughters, ABalap, had cell contact with the left-head precursor, ABplaa, based on our modeling results (Table S1), suggesting a specific signaling interaction between the two, which is consistent with previous cell-ablation results (Hutter and Schnabel 1995; Moskowitz and Rothman 1996). Notably, expression of the Notch ligand *lag-2* and the Notch receptor *lin-12* by LacZ-based transgenic assay suggested the signaling interaction at a later stage (i.e., between ABalapp and ABplaaa) (Moskowitz and Rothman 1996). However, our cell contact data suggest that ABalapa may play a bigger role than ABalapp in signaling the left head precursor (Fig. 4). The three cells stay in different z planes (Fig. 4A-C). Both daughters of ABalap express *apx-1*, but the relative contact area with ABplaaa is much greater for ABalapa (16.6%) than for ABalapp (5%) (Table S1). In addition, the daughters of ABalapa, but not those of ABalapp, are in contact with those of the daughters of the left head precursor (Movie S1), which further supports the more important role of ABalapa in signaling ABplaaa than ABalapp. These results suggest that the signaling effect in cell fate specification is achieved through consecutive signaling in multiple generations. It remains possible that two cells signal ABplaaa redundantly. Our reporter assay also showed that both Notch receptors may be redundantly involved in the signaling event, refining the previous finding that only a single ligand and receptor are involved in the third signaling event (Moskowitz and Rothman 1996).

**Fig 4.**
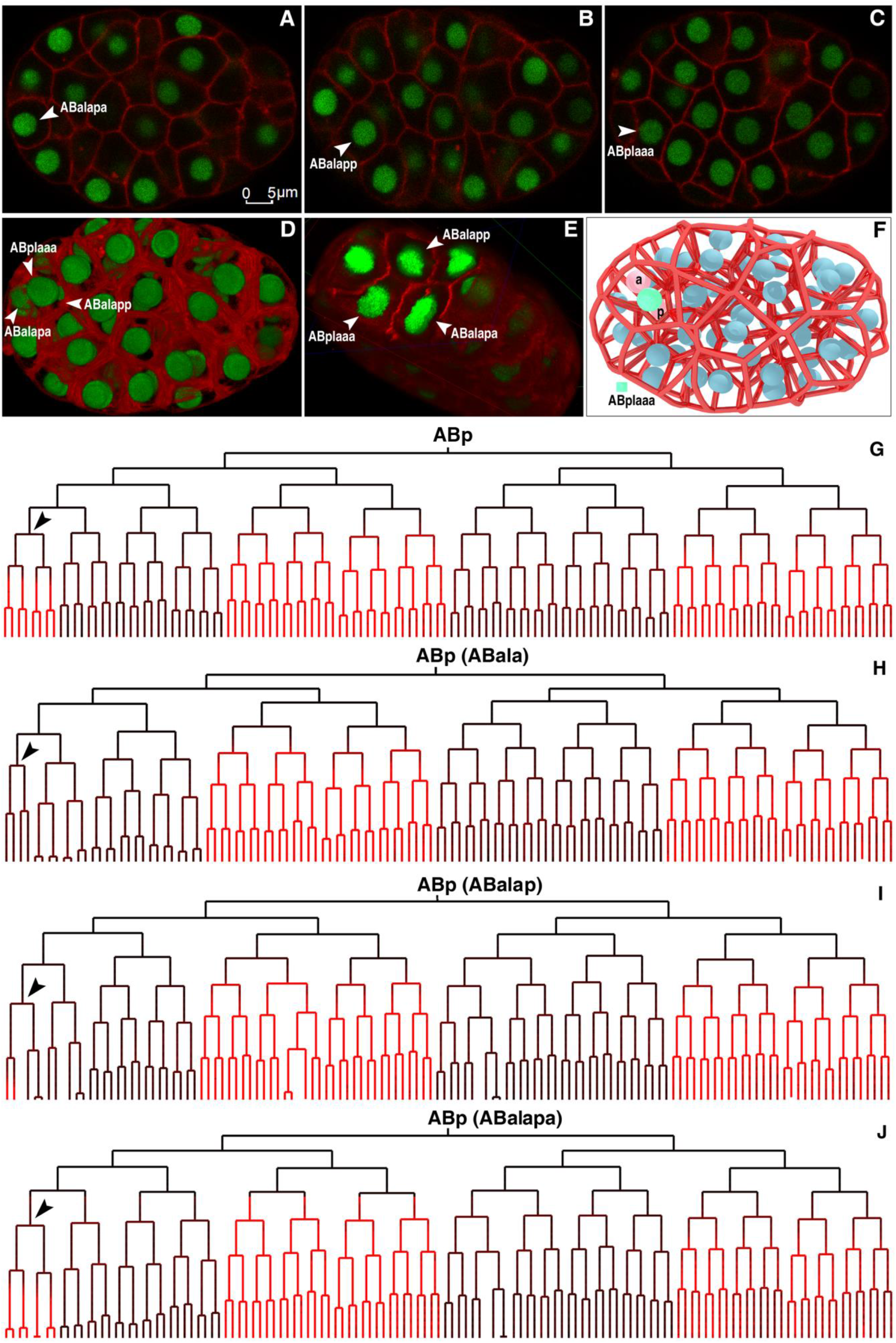
Refinement of 3^rd^ Notch signaling interaction in a 55-cell *C. elegans* embryo. A-C. Epifluorescence micrographs of different focal planes of the same *C. elegans* embryo (dorsal view with anterior to the left) focusing on cell ABplaaa (plane 26), ABalapa (plane 51) and ABalapp (plane 72), respectively, as indicated by arrowhead. Cell membranes and nuclei are colored in red and green respectively. D. 3D projection of epifluorescence micrographs. E. Cut-open view of the 3D projection in panel D showing cell boundaries. The projection is orientated to facilitate visualization of the cell boundaries. F. Modeling of cell boundaries in an embryo at approximately the same stage as in panel D. Nuclei of ABalapa and ABalapp are colored in red and indicated with an “a” and “p”, respectively, and the remaining nuclei colored in blue; ABplaaa nucleus is colored in green. G-J. Lineal expression of Pref-1::mCherry in “ABp” lineage of a wild-type embryo (G) or embryos with cell ablation (ablated cell indicated in parenthesis). Target cells of the 3^rd^ Notch signaling interaction are indicated with an arrowhead.

### Functional validation of the proposed cell pairs for the 3^rd^ Notch interaction

To experimentally validate the 3^rd^ Notch interaction, we first used cell membrane labeling coupled with cell lineage analysis (see Materials and Methods). Specifically, we performed 4D live-cell imaging of a *C. elegans* embryo ubiquitously expressing a nuclear and a membrane marker from the 4-cell stage up to the desired stage as estimated by wild-type lineaging trees (Ho *et al*. 2015). We then took a single 3D stack consisting of 110 focal planes for both GFP (nuclear) and mCherry (membrane) channels, which were rendered as a 3D projection (Fig. 4D). The 4D images allowed manual or automated tracing of cell identities, whereas the 3D projection permitted establishment of cell boundaries (Fig. 4 A-C). In agreement with our modeling results, the cell membrane labeling showed a higher confidence of contact with the left-head precursor by cell ABalapa than by cell ABplapp (Fig. 4D-E). The contact seems not obvious in modeled cell boundaries (Fig. 4F). This may be mainly due to the positional differences across the z axis.

We next verified whether the predicted signaling cell functions as expected using cell ablation technique. Given that *apx-1* is expressed in all ABala descendants, we decided to test whether the signaling interaction takes place in multiple generations as stated above by a combination of cell ablation and Notch target expression. We first ablated the cell ABala and examined the expression of a Notch target, *ref-1*, that is known to be expressed in the precursors of both the left and right heads (Neves and Priess 2005; Murray *et al*. 2012) (Fig. 4 G-H, Fig. S4A). As expected, ablation of the cell led to the specific loss of *ref-1* expression in the left-head precursor (Fig. 4H, Fig. S4A, 4A), demonstrating that the ligands expressed in this cell or its daughters are responsible for the signaling interaction. We next ablated the posterior daughter of ABala, ABalap, and re-examined the expression of *ref-1*. Interestingly, ablation of the cell abolished the *ref-1* expression in the posterior but not in the anterior descendants of ABplaaa (Fig. 4I and not shown), suggesting that some other signaling cell is responsible for the *ref-1* expression in the anterior descendants, or that the lost function in ABalap may be compensated by other ligand-expressing cells. We finally ablated the two daughters of ABalap (i.e., ABalapa and ABalapp). The former was proposed to be the signaling cell for ABlpaaa (Moskowitz and Rothman 1996), whereas our modeling and membrane labeling data supported a more important role for the latter in signaling ABplaaa (Fig. 4A-E). Unexpectedly, we observed that the *ref-1* expression in ABplaaa descendants after either ablation was comparable to that of the wild type (Fig. 4J and data not shown, Figs. S4, S5C-E, S6). Taken together, our results suggest that the induction of left-head specification is achieved by a multiple round of signaling from consecutive cell cycles, which is especially true during the late stage of embryogenesis. The results also suggest redundant features of Notch signaling in regulating fate specification.

### Identification of the proposed cell pairs for the 4^th^ Notch signaling in *C. elegans* embryo

Previous studies suggested that one or both of the MSap daughters are the signaling cell(s) for fate specification of ABplpapp, the great-grandparent of the excretory cell (a functional equivalent of the human kidney), but the exact identities of the signaling cells remain elusive (Moskowitz and Rothman 1996; Priess 2005). To establish the identity of the signaling cell, we first examined our modeling results on cell contact, which suggest that only one of the MSap daughters (i.e, MSapp but not MSapa) is in contact with the excretory cell precursor (Fig. 5, Table S1). Consistent with this, a 3D projection of labeled cell membranes showed that it is MSapp but not MSapa that is in contact with the ABplpapp cell (Fig. 5A-D, Movie S2). To further validate the interaction between the two cells, we examined the lineal expression of both Notch ligands and receptors. We observed that one Notch receptor, *lin-12*, was expressed in all descendants of ABplp, the great-grandparent of ABplpapp (Fig. 3D). Consistent with our modeling results, the GFP reporter of one Notch ligand, *lag-2*, was specifically expressed in MSapp but not in MSapa (Fig. 5E, Fig. S3 D-E, I-J), further supporting that MSapp is the signaling cell for ABplpapp. Notably, one daughter of MSapp, MSappa, was also in contact with ABplpapp, indicating that the signaling interaction is further relayed in the next cell cycle.

**Fig 5.**
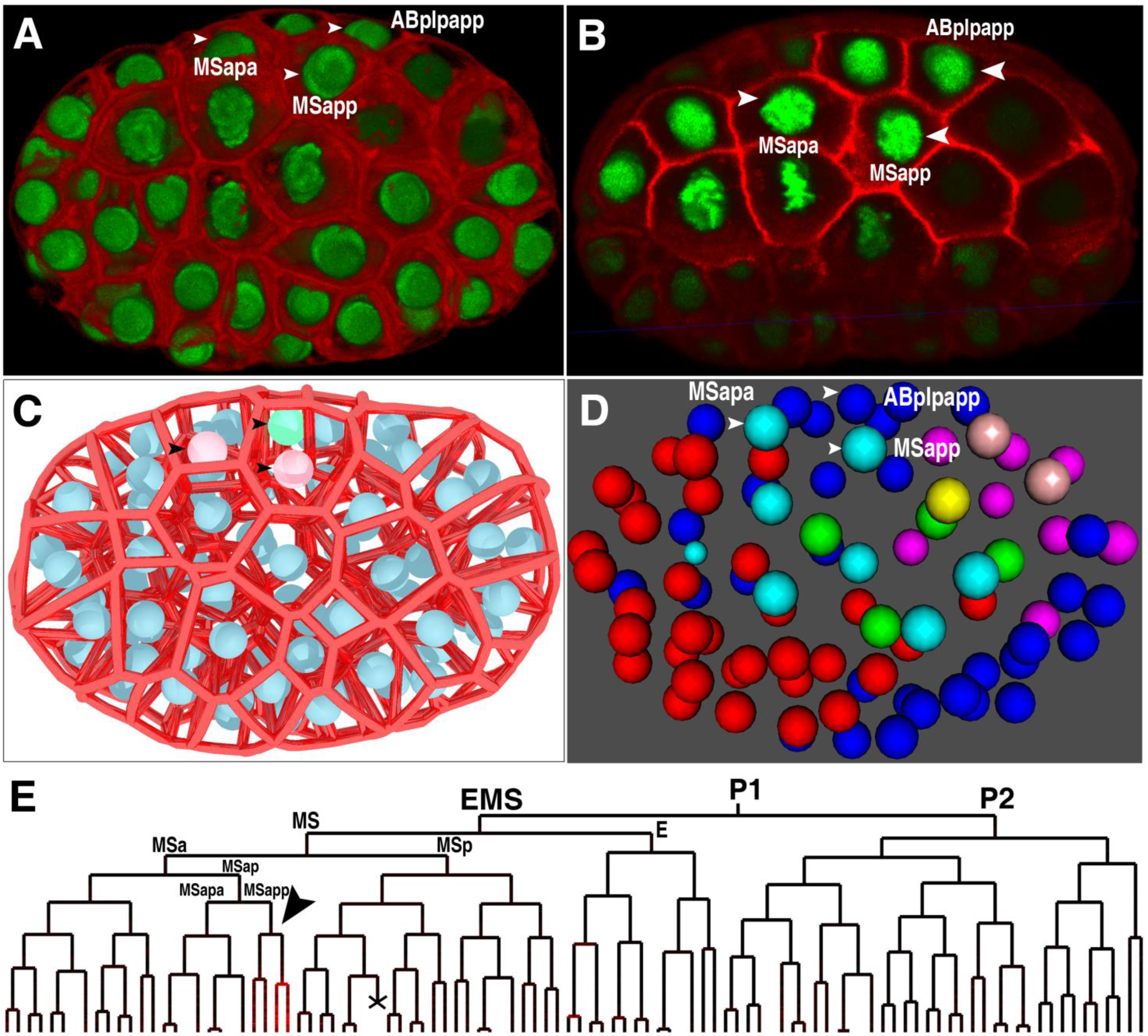
Refined 4^th^ Notch signaling interaction in *C. elegans* embryo. A. Shown is a 3D projection of epifluorescence micrograph of an 87-cell embryo with cell membranes labelled by mCherry (red) and nuclei by GFP. One or both of MSap daughters were previously proposed to signal excretory cell precursor, ABplpapp. The three cells are indicated with an arrowhead. B. Cut-open view of the same embryo as in panel A showing cell boundaries. The embryo is oriented so that the boundaries of interest are most obvious. C. Modeling of cell boundaries of the same embryo as in panel B with the same three cells indicated with an arrowhead. ABplpapp is colored in green and two MSap daughters in red and the remaining nuclei in blue. D. 3D space-filling model of an embryo at the same stage as in panel A. Red: ABa; dark blue: ABp; light blue: MS; green: E; pink: C; brown: D; yellow: P4. The same three cells as in panel A are indicated with an arrowhead. E. Lineal expression of a Notch ligand, *lag-2*, in MSapp (red) indicated with an arrowhead. Cell death is indicated with an “X”.

### Evidences of Notch signaling in later AB descendants

The transcription factor *pal-1* is expressed in ABplppppp, the grandparent of the anal depressor muscle and an intestinal muscle, and appears to be a direct target of Notch signaling required for rectal development (Edgar *et al*. 2001). The signaling cells for this interaction appear to be descendants of MSapa or MSapp (Priess 2005), but the exact identities of the signaling cells remain elusive. Our modeling results predicted a reproducible cell contact between MSappp and ABplpppp, the parent of ABplppppp (Fig. 6D, Table S1). Cell membrane labeling and a space-filling model support the contact between the two cells (Fig. 6A-D), but not between MSapa daughters and ABplpppp (Table S1), demonstrating that MSappp is more likely to be the signaling cell for ABplpppp that is required for *pal-1* expression in ABplppppp.

**Fig 6.**
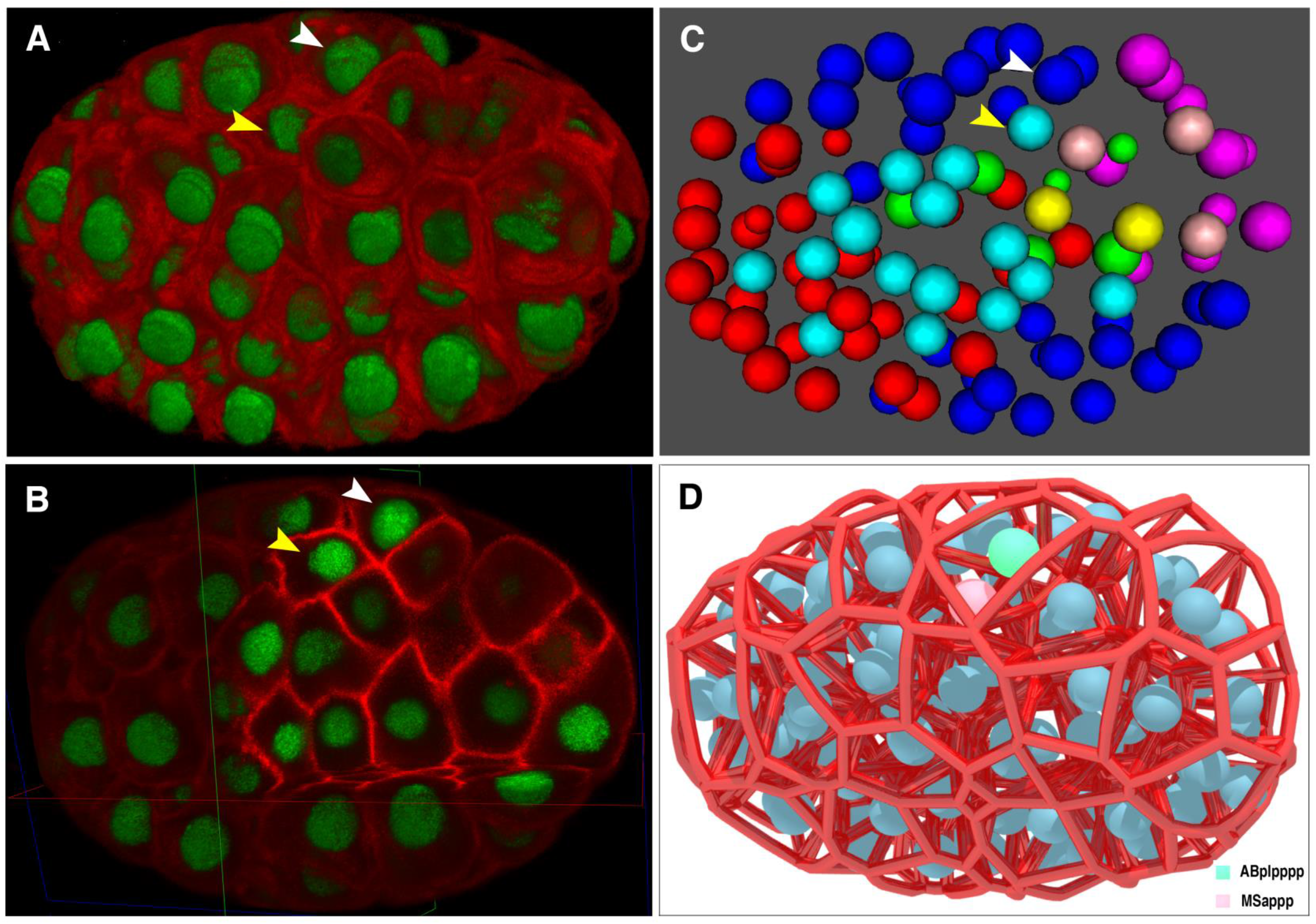
Identities of cells for fifth Notch signaling interaction in *C. elegans* embryo. A. 3D projection of a 96-cell embryo with cell membranes and nuclei colored in red and green, respectively. Signaling interaction was proposed to take place between two cells, MSappp (yellow arrowhead) and ABplpppp (white arrow head). B. Cut-open view of the same embryo as in panel A showing cell boundaries. C. 3D space-filling model of a 96-cell *C. elegans* embryo with cell pairs similarly color coded as in panel A. D. Modeling of cell boundaries in a 96-cell *C. elegans* embryo. Nuclei of MSappp and ABplpppp are colored in red and green, respectively and the remaining nuclei in blue.

In a wild-type embryo of approximately the 300-cell stage, a contact between two bilaterally symmetric AB descendants, ABplpapppp and ABprpapppp, appears to be required for a Notch interaction for the former to develop into a neuron and a rectal epithelial cell (Bowerman *et al*. 1992). Our modeling results predicted a contact between the two cells with a high level of confidence (Table S1). Lineal expression of a Notch receptor, *lin-12*, was observed in ABplpapppp (Fig. 3D) although that expression of both of our Notch ligands was not observed in ABprpapppp, suggesting other Notch ligands may be involved in the interaction.

### A web-based utility for access to the cell-cell contact data over *C. elegans* embryogenesis

To facilitate the intuitive use of our cell contact map, we developed a webpage that allows online query and navigation of cell contacts over embryogenesis (Fig. S7). One can access the contacts relevant to their cell of interest by searching for the cell name or by navigating through a lineage tree. The output will show all cells that are in contact with the cell of interest in a graphical representation in which the thickness of the bars is proportional to the predicted score of a specific contact. The website is accessible through the link: http://ccccm.bionetworks.ml/.

## Discussion

Signaling interaction plays a key role in breaking of division symmetry during metazoan development. Accurate and systematic identification of the interactions at cellular resolution during development is critical for understanding molecular mechanism of symmetry breaking but is technically challenging (Zacharias *et al*. 2015). This is especially true during a late proliferative stage of embryogenesis due to the difficulties in establishing contacting cells and their identities (Bao *et al*. 2006; Richards *et al*. 2013). It is also challenging to generate the native expression dynamics of signaling molecules at cellular resolution for each cell cycle.

Here, we present an automated platform that allows accurate identification of signaling interactions at cellular resolution during the proliferative stage of *C. elegans* embryogenesis. This was achieved by a combination of computer modeling of cell contact, automated cell lineaging and single-cell gene expression profiling. The cell contact map calibrated with both membrane labeling and known signaling interactions lays a foundation for systematic identification of signaling interactions. Applying the platform in *C. elegans* not only allows validation and refinement of the existing Notch signaling interactions but also permits identification of multiple novel signaling interactions especially during a relatively late embryonic stage. The method can be applied to the characterization of any other signaling interactions. It should be noted that the Voronoi modeling is an approximation of cell surface. Predicted interactions with a smaller surface area in contact may or may not be functionally relevant. Alternatively, some functional contacts might be missed out in our list due the empirical cutoff we used in the modeling process. Functional test is required for making a functional calling of a functional contact.

Although many existing fate specifications were proposed to be triggered by a single signaling event, our analyses suggest that fate specification may depend on multiple signaling interactions that take place consecutively across cell divisions. For example, though our cell contact data and membrane labeling results support that it is ABalapa that mediates the third Notch interaction (Fig. 4) instead of ABalapp as described previously (Moskowitz and Rothman 1996), ablation of either ABalapa or ABalapp doesn’t affect *ref-1* expression in ABplaaa descendants. We propose that the relay of signaling interactions over multiple generations may be a common practice for breaking of division symmetry as suggested earlier based on lineal expression of Wnt components (Zacharias *et al*. 2015). Alternatively, the interaction might be very brief. Once the signaling cell is born the signaling event might happen very quickly and ablation of that cell soon after its birth might not be enough to block the signaling interaction. We also observed frequent redundancy of signaling interactions which may serve to increase the robustness of a developmental process.

All of the expression patterns for Notch ligands and receptors are derived from a fusion between their promoter sequences and GFP with a heterogeneous 3’ UTR from *his-72*. Therefore, these vectors may capture only the zygotic but not maternal expression (Murray *et al*. 2008). In addition, the arbitrarily chosen fragment may not necessarily contain all of the functional elements required to drive its native expression. Because all of the expression patterns are derived from a single-copy transgene, some of them may be too dim to be detectable. Therefore, certain expressing cells or stages may be missing in our dataset. For example, the expression of *lag-2* was seen in ABala descendants by extrachromosomal array (Moskowitz and Rothman 1996), but not in our transgenic strain (Fig. S3 D-E), which could be because the expression driven by a single-copy transgene is too dim to be detected or because some *cis*-elements are lacking in the promoter used. Use of a brighter reporter, for example, Ruby3 (Bajar *et al*. 2016), may facilitate the visualization of single-copy transgenes. In summary, we present a new map of cell-cell contacts in *C. elegans* embryogenesis. We applied the map together with 4D imaging-based cell lineage analysis to refine previously described cell inductions. We finally develop a website that potentially becomes a valuable resource to the *C. elegans* community for intuitive and easy access to cell-cell contacts.

## Materials and Methods

### Modeling of cell-cell contact

Prediction of cell surface is performed using the Voronoi segmentation algorithm (Franz Aurenhammer 1991; Atsuyuki Okabe, Barry Boots 2000) with the “Voro++” library(Rycroft 2009) using the output from StarryNite as an input, which contains 3D coordinates for all nuclei at a 1.5-minute interval from 4 to 350 cells of a *C. elegans* embryo. One caveat of the method is the segmentation of the cells located at the edge of an embryo, where a false positive cell contact may be predicted as reported previously (Hench *et al*. 2009). To solve this issue, for each embryo, a 3D convex hull was generated as a proxy for embryo boundary. Given the reproducible migration of cells at the embryo boundary, cell surface and contact areas were computed with cells’ coordinates and the 3D convex hull with “Voro++”.

3D coordinates from 91 wild-type *C. elegans* embryos were individually modeled to define cell contacts for each time point (1.5 minute) for all embryos. To evaluate the variability of cell contacts among embryos, cell contact areas were compared against each embryo using “cell stage”, i.e., the number of cells in a given embryo, rather than the absolute developmental time. This would minimize the complications associated with variability in developmental timing.

### Visualization of cell boundary at desired stage

A strain ZZY0535 was made by crossing the lineaging strain RW10029 expressing GFP lineaging markers with strain OD84 expressing a membrane marker, P*pie-1*::mCherry::PH (PLC1delta1) (see Table S3). The three markers were rendered triply homozygous.

For visualization of cell contact by fluorescence membrane labeling, a 4-cell embryo with desired developmental timing was selected for 4D imaging with a Leica SP5 confocal microscope using the similar settings as those used for automated lineaging till the embryo developed to the desired stage. Timing for presence of a cell of interest was estimated based on our lineaging results of the 91 wild-type embryos(Ho *et al*. 2015). Imaging with live data mode was switched to normal mode to take a single stack consisting of 110 focal planes with suitable AOTF compensation using a pinhole of 1.6 AU and three-line accumulation. Images were acquired from both GFP and mCherry channels. Identity of the cell of interest was resolved by manually navigating through the image stacks using Leica Application Suite X (LAS X). The 3D stack of the embryo was used to reconstruct the 3D volume projection with LAS X. The embryo was rotated to a proper orientation to facilitate visualization of the desired cell boundary. Part of the embryo was cut open across different axes for visualization of contact.

### Cell ablation coupled with cell lineage analysis

For cell ablation coupled with automated lineaging, a 4-cell embryo with desired orientation was selected for 4D-imaging till the cell targeted for ablation was born. Manual tracing of the targeting cell was performed with the help of the lineaging markers. Immediately after the target cell completed mitosis, imaging was terminated and the following procedures were performed within 1.5 minutes: switch the imaging mode from live data mode to normal mode; focus on the middle plane of the target cell nucleus by fine-tuning the Z-Galvo; select the bleaching point from the panel and create a region of interest (ROI) in the middle of the target cell nucleus in a preview panel; turn off all the fluorescence detectors except the one for the DIC and switch the filter to “Substrate”; set the bleaching time (40 seconds for ABala, 20 seconds for all others); temporally close the shutters for all the wavelengths except the pulsed diode laser (PDL 800-B, PicoQuant), which emits 405nm laser beam; tune it to 100% intensity and start the bleaching; and once completed, switch back to the live data mode and resume the 4D-imaging as usual.

### Generation of single-copy transgenic lines

Single-copy transgene consisting of a fusion between the promoter of a Notch ligand/receptor and GFP was generated using miniMos technique(Frokjaer-Jensen *et al*. 2014). The primer sequences used for amplifying the promoter sequences were listed in Table S4. The miniMos targeting vector pCFJ909 was modified to include a genomic coding region of *his-24* upstream of the GFP coding sequence to facilitate nuclear localization for automated lineal expression profiling as described(Zhao *et al*. 2010c). Multiple independent strains were produced for each promoter. A single strain that shows expression patterns consistent with the remaining ones was genotyped by inverse PCR and crossed with the lineaging strain RW10226. Both lineaging and Notch markers were rendered homozygous for automated lineaging and lineal expression profiling as described(Zhao *et al*. 2010c).

### 4D live cell imaging, automated lineaging and profiling of lineal gene expression

Imaging was performed in the similar way to that described previously(Shao *et al*. 2013). Briefly, lineaging strain, RW10226(Zhao *et al*. 2010a), ubiquitously expressing nuclear mCherry, was crossed with the strain expressing a fusion between the promoter of a Notch component and GFP. Both the lineaging markers and the promoter fusion were rendered homozygous before lineaging. 4D imaging stacks (roughly 0.7 µm/stack) were sequentially collected for both GFP and RFP (mCherry) channels at a 1.5-minute interval for a total of 240 time points using a Leica SP5 confocal microscope as described(Shao *et al*. 2013). Automated profiling of lineal expression was performed as described(Zhao *et al*. 2010c).

### Worm strains and maintenance

All the animals were maintained on NGM plates seeded with OP50 at room temperature unless stated otherwise. The genotypes of the strains used in this paper were listed in Table S3.

## Acknowledgements

We thank Mr. WS Chung for logistic support and helpful discussion with the members of Z Zhao’s laboratory. This work is supported by the Hong Kong Research Grants Council (Project numbers HKBU5/CRF/11G, HKBU263512 and HKBU12103314) and HKBU Faculty Research Fund (FRG2/13-14/063), HKBU Strategic Development Fund for Environmental Genetics and the National Natural Science Foundation of China (Project 61702396) and the China Postdoctoral Science Foundation (Project 2016M600769). Some of the strains used in this study were provided by *C. elegans* Genetic Center, which is funded by NIH Office of Research Infrastructure Programs (P40 OD010440).

## Author contributions

L.C modeled the cell contacts and H.C.K.N contributed to the dataset. M.K.W generated 3D projections of membrane-labeling and performed cell ablation. V.W.S.H, L.Y.C and X.R made the transgenic strains and produced lineal expressions. H.Y and Z.Z conceived the project. L.C and Z.Z wrote the manuscript.

## Supporting Information Legends

### Supporting Tables

Table S1. List of cell pairs between which an effective cell contact is called.

Table S2. List of cells expressing Notch ligands or receptors.

Table S3. List of strains and its genotypes

Table S4. List of PCR primers for amplification of promoters for Notch ligands and receptors

### Supporting Figures

Figure S1. Occurrence distribution of the ratio of modeled contact area between MS and ABalp (A) or ABara (B) relative to average total cell surface area at the current time point. Second Notch signaling interactions are well established between the two cell pairs.

Figure S2. Comparison of brood sizes between unmounted (non-pressurized control) and mounted (mounting under pressure) embryos. Shown are boxplots of brood sizes that were scored from eight adults for each group with the total number of counted embryos indicated.

Figure S3. Lineal and spatial expression of Notch components up to 350 cells. A-C. Lineal expression of two Notch receptors, *glp-1* (A) and *lin-12* (B), and a Notch ligand, *apx-1* (C), in P1 sublineage. D-E. Lineal expression of a Notch ligand, *lag-2*, in ABa (D) and ABp (E) sublineage. F-I. Space-filling models showing spatial expression of the above four genes in a 350-cell *C. elegans* embryo. Brightness in red corresponds to expression intensity. J. Spatial expression of *lag-2* based on their lineal origins in a 350-cell *C. elegans* embryo. ABalaappaa, red; ABalapapaa, blue; ABalppaaaa, pink; ABalppaapa, yellow; ABalappaaa, green; MSapp descendants, cyan.

Figure S4. Lineal expression of *ref-1* in ABa sublineage of embryos before (A) or after (B-E) cell ablation. Names of the ablated cells are indicated in parenthesis with approximate ablation timing indicated by an arrow. Note a loss of *ref-1* expression in descendants of the ablated cells.

Figure S5. Lineal expression of *ref-1* in P1 sublineage of embryos before (A) or after (B-E) cell ablation. Programmed cell deaths are indicated with arrow.

Figure S6. Lineal expression of *ref-1* in ABa (A) or ABp (B) sublineages of embryos before (A) or after (B-E) ablation of ABalapp as indicated on the top of each lineage tree. Cell ablation is indicated by “arrowhead” and the left head precursor, ABplaaa, is indicated by “arrow”.

Figure S7. Screenshot of the output by searching website of *C. elegans* Cell-Cell Contact Map (CCCCM) using cell “ABplaaa” as a query. The querying cell is highlighted in red while its contacting cells in blue in both lineage tree (top, up to 350 cells) and network schematics (bottom). Relative contact area is shown in proportional to the thickness of the bar connecting contacting cells. Details of cell contact information are shown on the bottom right. Only contacts that satisfy our threshold are shown. Cells can also be queried using by navigating lineage tree as shown in bottom left.

### Supporting Movies

Movie 1. A time-lapse movie showing the contacts between ABplaaa and ABalpap and those between their daughters.

Movie 2. A time-lapse movie showing the contacts between excretory cell precursor, ABplpapp and MSapp.

